# Ultra-High Resolution TR-external EPIK with Deep Learning Image Reconstruction for Enhanced Characterisation of Cortical Depth-dependent Neural Activity

**DOI:** 10.1101/2025.02.21.639447

**Authors:** Seong Dae Yun, Jaejin Cho, Patricia Pais-Roldán, N. Jon Shah

**Author notes:** **To whom correspondence may be addressed:** Dr. Seong Dae Yun, Institute of Neuroscience and Medicine 4, Forschungszentrum Jülich, 52425 Jülich, Germany. Seong Dae Yun and Jaejin Cho contributed equally to this work.

## Abstract

The detection of neural signals using functional MRI at the laminar or columnar level enables non-invasive exploration of fundamental brain function processing and the interconnected pathways within intracortical tissues. The growing interest in this research area is driven by advancements in fMRI acquisition techniques that enhance spatial resolution for high-fidelity mapping. However, the submillimetre voxel sizes commonly employed in layer-specific fMRI studies raise concerns about relatively low signal-to-noise ratios and increased image artefacts in reconstructed images, ultimately limiting the precise delineation of cortical depth-dependent functional activities.

This work aims to address this issue by incorporating a deep learning technique for enhanced image reconstruction of the submillimetre fMRI data, acquired with echo-planar-imaging with keyhole (EPIK) combined with the repetition-time-external (TR-external) EPI phase correction scheme. Our network was trained in a self-supervised, scan-specific manner using the sampling strategy from zero-shot self-supervised learning (ZS-SSL) method, which has gained attention for high-resolution MR image reconstruction.

Healthy volunteers participated in this study, and the performance of the developed method was evaluated in direct comparison to the conventional reconstruction method using datasets acquired at 7T. The deep learning reconstruction produced reconstructed images with significantly higher SNR than the conventional method, which was further quantitatively validated through histogram analysis. This enhancement was consistent across all slice locations, demonstrating the reliability of the scan-specific deep learning technique.

## 1. Introduction

The investigation of cortical depth-dependent functional activity has gained increasing popularity in the neuroimaging community, as it offers deeper insights into the laminar organisation of the cortex, which is crucial for understanding how the brain processes information. The unique advantages and continuous developments of functional MRI (fMRI) over other neuroimaging modalities, particularly its unrivalled in vivo spatiotemporal resolution and non-invasiveness (Yun et al., 2024a), have led to its widespread use in layer-specific fMRI studies. For the successful delineation of layer-specific functional profiles, fMRI techniques have been advanced with a focus on the detection of signals originating from actual activation sites and their depiction in the spatial representation of the brain with a high mapping fidelity (Yun et al., 2024b).

One widely adopted methodological approach to improve the mapping specificity is to increase the spatial resolution of fMRI acquisition. Given the fact that the human cortex is approximately 3-5 mm thick in which six layers are organised, it is highly demanded to achieve a submillimetre spatial resolution, and most previous layer-specific fMRI studies have employed a nominal spatial resolution of 0.5 ∼ 0.9 mm (Beckett et al., 2020; Berman et al., 2021; Han et al., 2022; Iyyappan et al., 2024; Kashyap et al., 2021; Kay et al., 2019; Yun et al., 2022. While this resolution effectively resolves cortical layer-specific activation, it comes at the cost of low signal-to-noise ratio (SNR) due to the intrinsically smaller voxel sizes.

This concern has predominantly been addressed by using ultra-high field strengths in the studies. However, accelerated imaging techniques, such as parallel imaging, partial Fourier and multi-band techniques (Margosian et al., 2007; Ra and Rim 1993; Setsompop et al., 2011; Weaver 1988) commonly employed in submillimetre fMRI acquisition using EPI-based methods, also contribute to further SNR degradation. Moreover, the use of these acceleration techniques introduces additional image artefacts, such as aliasing and blurring during, the reconstruction process. These artefacts become more pronounced at higher acceleration factors to increase the spatial resolution, where their complete elimination is limited to some extent with the standard reconstruction approach. The relatively low SNR and poor image quality are major factors that limit the reliable and reproducible detection of functional responses. This issue can be mitigated using de-noising techniques, as demonstrated in previous studies (Carone et al., 2017; Knudsen et al., 2023). However, these techniques need to be tailored to the specific noise distribution patterns, and may introduce concerns about spatial smoothing, which can result in potential loss or blurring of functional signals (Hoeppli et al., 2023).

Recently, deep learning has demonstrated significant potential in MR image processing, particularly in enhancing the reconstruction of high-resolution MR data under relatively low SNR conditions (Aggarwal et al., 2018; Desai et al., 2023; Hammernik et al., 2018). In deep learning, features and representations are automatically extracted through the training of large datasets, typically performed in a data-driven manner. By learning from various noise patterns and image artefacts, deep learning techniques can effectively eliminate these artefacts while preserving the fine-scale signals. However, the application of deep learning to layer fMRI remains in its early stages, primarily due to the need for new architectures, as traditional deep learning models may not be well-suited for the unique structure of the ultra-high resolution fMRI data. The large size of these high-resolution datasets demands extensive computational resources for training, further increasing the challenge.

Recently, self-supervised learning techniques have gained attention for high-resolution MR image reconstruction, with zero-shot self-supervised learning (ZS-SSL) methods showing comparable reconstruction quality with conventional supervised approaches (Yaman et al., 2020a and 2020b). This is particularly valuable for EPI, where rapid sampling, susceptibility to artefacts, and limited availability of high-fidelity ground-truth data make traditional supervised methods challenging to implement. By learning directly from the data within a single scan, zero-shot self-supervised approaches mitigate the need for large training sets while achieving robust EPI reconstructions. Moreover, they have been extended successfully to other MRI applications, including quantitative imaging and diffusion imaging (Cho et al., 2023; Jun et al., 2023).This work primarily focuses on the development of a self-supervised deep learning network to enhance image reconstruction quality in ultra-high-resolution fMRI data. Specifically, the fMRI protocol was configured using echo-planar imaging with keyhole (Caldeira et al., 2019; Küppers et al., 2022; Pais-Roldán et al., 2023; Shah & Zilles, 2003, 2004; Yun et al., 2013, 2019, Yun & Shah, 2017; Zaitsev et al., 2001, 2005) combined with TR-external EPI phase correction (TR-external EPIK)(Yun et al., 2022), which has demonstrated superior spatial resolution and brain coverage, while maintaining comparable functional detection performance to the community standard method, EPI. The developed scan-specific deep learning image reconstruction technique is applied to submillimetre TR-external EPIK data (0.875 × 0.875 mm^2^), and its performance is evaluated in comparison to the conventional reconstruction method.

## 2. Materials and Methods

### 2.1. Study subject and ethics statement

Healthy volunteers were recruited and underwent standard safety screening procedures. None had any medical, neurological, or psychiatric conditions. After receiving a full description of the study, written informed consent was obtained prior to scanning. The study protocol, along with the screening questionnaires and consent forms, was approved by the local institutional review board at RWTH Aachen University, Germany (EK 346/17).

### 2.2. Ultra-high resolution TR-external EPIK

Figure 1 shows a schematic representation of the TR-external EPIK sequence diagram (Yun et al., 2022). The navigator echoes, used for the correction of N/2 ghost artefacts, are acquired in a separate excitation loop preceding the main acquisition kernel, allowing for more phase-encoding lines to be accommodated at a given TE for a higher spatial resolution. The flip angle (α_PC_) and TR (TR_PC_) in this excitation loop are set much smaller than those in the main excitation loop, minimising the increase of additional radio-frequency energy accumulation and acquisition time.

**Figure 1.**
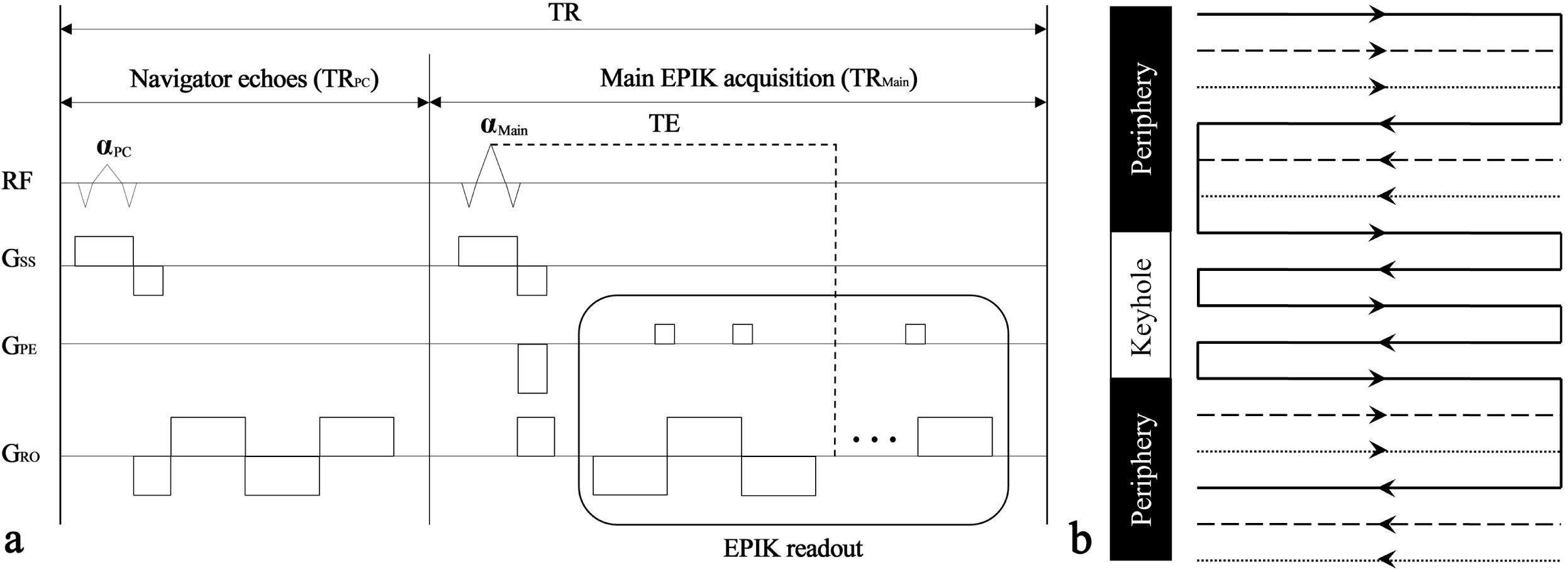
Schematic representation of the TR-external EPIK sequence diagram. (**a**) TR-external phase-correction scheme where three navigator echoes are acquired in a separate excitation loop, while main data are acquired in the subsequent excitation loop using EPIK readout. (**b**) Three-shot EPIK acquisition scheme, where the central k-space region (denoted as the keyhole) is fully sampled at every temporal scan while the peripheral k-space regions are sparsely sampled resembling three-shot EPI. The solid, dashed and fine-dashed lines in the peripheral regions indicate the sampling positions performed at the 1st, 2nd and 3rd measurements, respectively.

Data from healthy volunteers were acquired with the following imaging parameters: TR = 3500 ms (i.e., 12 ms for TR_PC_ and 3488 ms for TR_Main_), TE = 220ms, FOV = 2100×0210 mm^2^, matrix = 2880×0288 (0.730×00.730mm^2^), slices = 48 with 1 mm thickness and a 2 mm gap, partial Fourier = 7/8, parallel imaging = 5-fold, multi-band = none, and α_PC_/α_Main_ = 9°/90°. The same imaging parameters were repeated with the phase-encoding direction reversed to correct for geometric distortions. The above configuration was applied on a Siemens Magnetom Terra 7T scanner equipped with a single-channel Tx/32-channel Rx Nova medical coil provided by the manufacturer.

### 2.3. Image reconstruction using the deep learning network

Figure 2a illustrates the proposed network architecture for ultra-high-resolution fMRI data. A data consistency (DC) layer enforces agreement between the acquired raw data and the reconstructed images, while two unrolled CNN denoisers operate in both *k*-space and image-space. Previous studies have demonstrated that incorporating denoisers in both domains improves performance metrics such as peak SNR (PSNR) and structural similarity index measure (SSIM) (Aggarwal et al., 2019; Eo et al., 2018). The network also employs the virtual coil (VC) technique (Blaimer et al., 200), which generates additional *k*-space information by leveraging conjugate symmetry from actual coil data. This extra information is particularly beneficial for partial Fourier acquisitions, further enhancing reconstruction quality.

**Figure 2.**
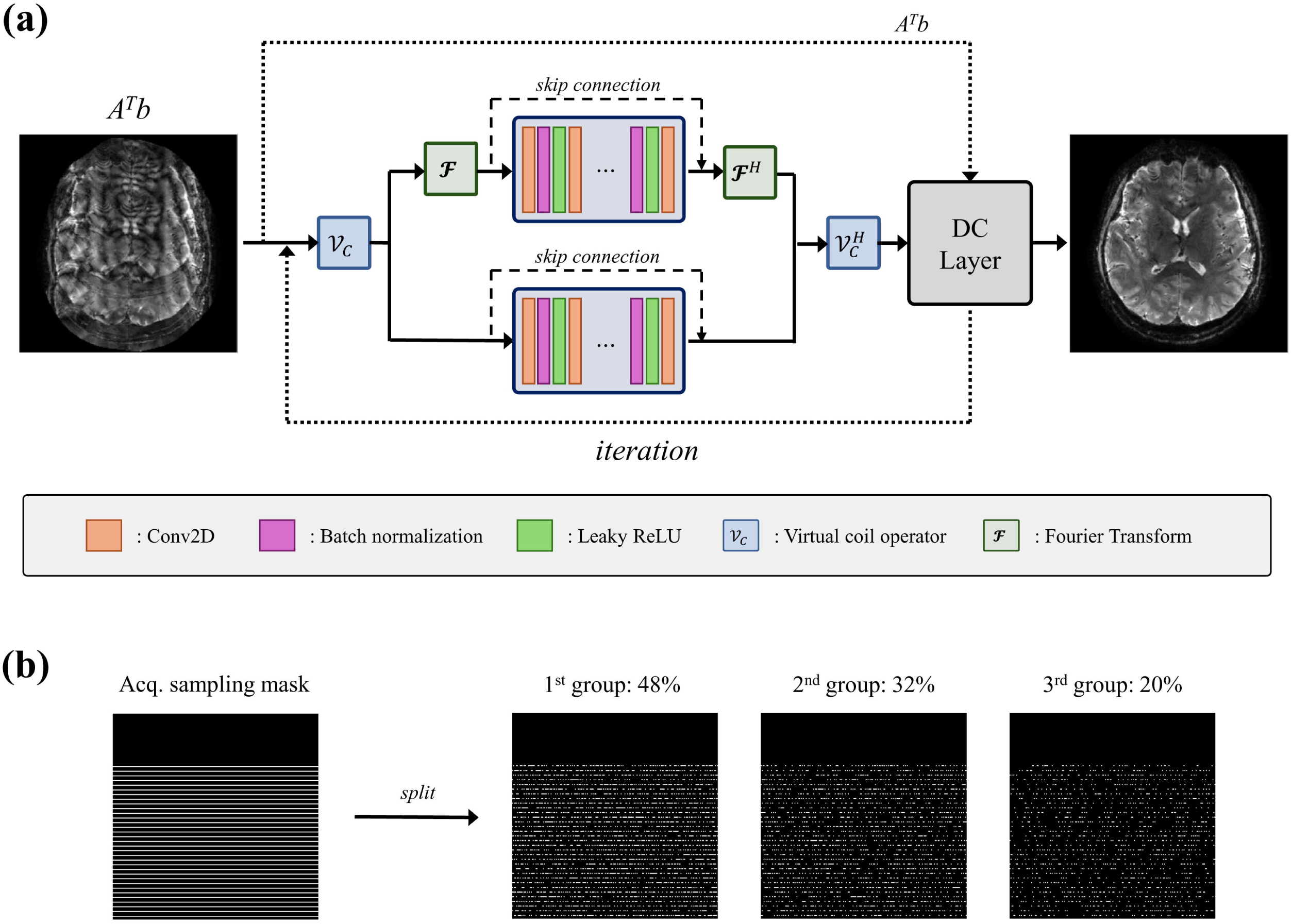
Schematic representation of the proposed deep learning network. (**a**) The proposed network diagram for ultra-high-resolution fMRI. (**b**) Sampling splits for self-supervised learning.

To train our network in a self-supervised, scan-specific fashion, we adopt the sampling splitting strategy proposed in ZS-SSL (Yaman et al., 2020a). Specifically, we divide the sampling mask of each scan into three groups with a ratio of 0.48:0.32:0.20, as shown in Figure 2b. During the training phase, k-space data in the first sampling group provides input to the network, while the training loss is computed over the second group. We create 20 distinct sets for the first two groups per slice to facilitate the training. In the validation phase, the network input is formed by combining the first two groups, and the validation loss is then measured using the third group. This procedure identifies the optimal network model and determines the early stopping point. Finally, during inference, all sampling points are used to generate the best possible reconstruction. For robustness, we trained only a single network across all slices and all time points.

All neural network implementation were conducted with Python, using the Keras library in TensorFlow 2.11.1. NVIDIA RTX A6000 (RAM: 48 GB) was used for training, validating, and testing the network. Each denoising CNN in k-space and image-domain consists of 16 layers of which the depth is 38. The filter size was 3 × 3, and the number of trainable parameters was 396,578. The number of iterations was 8. For training the network, we employed the Adam optimizer with a learning rate of 1e-3. Leaky ReLU was used as the activation function. In this study, we used sum of the normalized root mean square error (NRMSE) and normalized mean absolute error (NMAE) as the loss functions.

### 2.4. Qualitative and quantitative evaluation of reconstructed image quality

The quality of reconstructed images obtained from the developed deep learning techniques was assessed in direct comparison to that from the conventional method. The potential improvement in SNR of the deep learning-reconstructed images was evaluated by examining the dispersion of histogram distributions for signals within the grey matter (GM) and white matter (WM) regions. These regions of interest (ROIs) were defined using the standard SPM12 segmentation routine (Wellcome Department of Imaging Neuroscience, UCL, London, UK). For each ROI, the mean ± SD of each ROI of the signals was computed for both the conventional and deep learning reconstructed results.

## 3. Results

### 3.1. Reconstructed images

Figure 3a shows reconstructed images obtained from both conventional and deep learning reconstructed methods at four representative slice locations. The results from the conventional method exhibit noticeable noise contamination mostly due to the use of a relatively high parallel imaging acceleration factor. However, the noise is significantly reduced in the deep learning reconstructed results across all slice locations presented in the figure. This improvement in SNR is more clearly confirmed in the enlarged view of the selected rectangular ROIs, as shown in Fig. 3b. This figure highlights the effective reduction of noise contamination arising from the acceleration technique, with no significant visible loss of spatial resolution. Figure 4 displays the sagittal and coronal representation of the reconstructed axial slices, demonstrating the reliable performance of the developed deep-learning reconstruction method in improving SNR across all slice locations.

**Figure 3.**
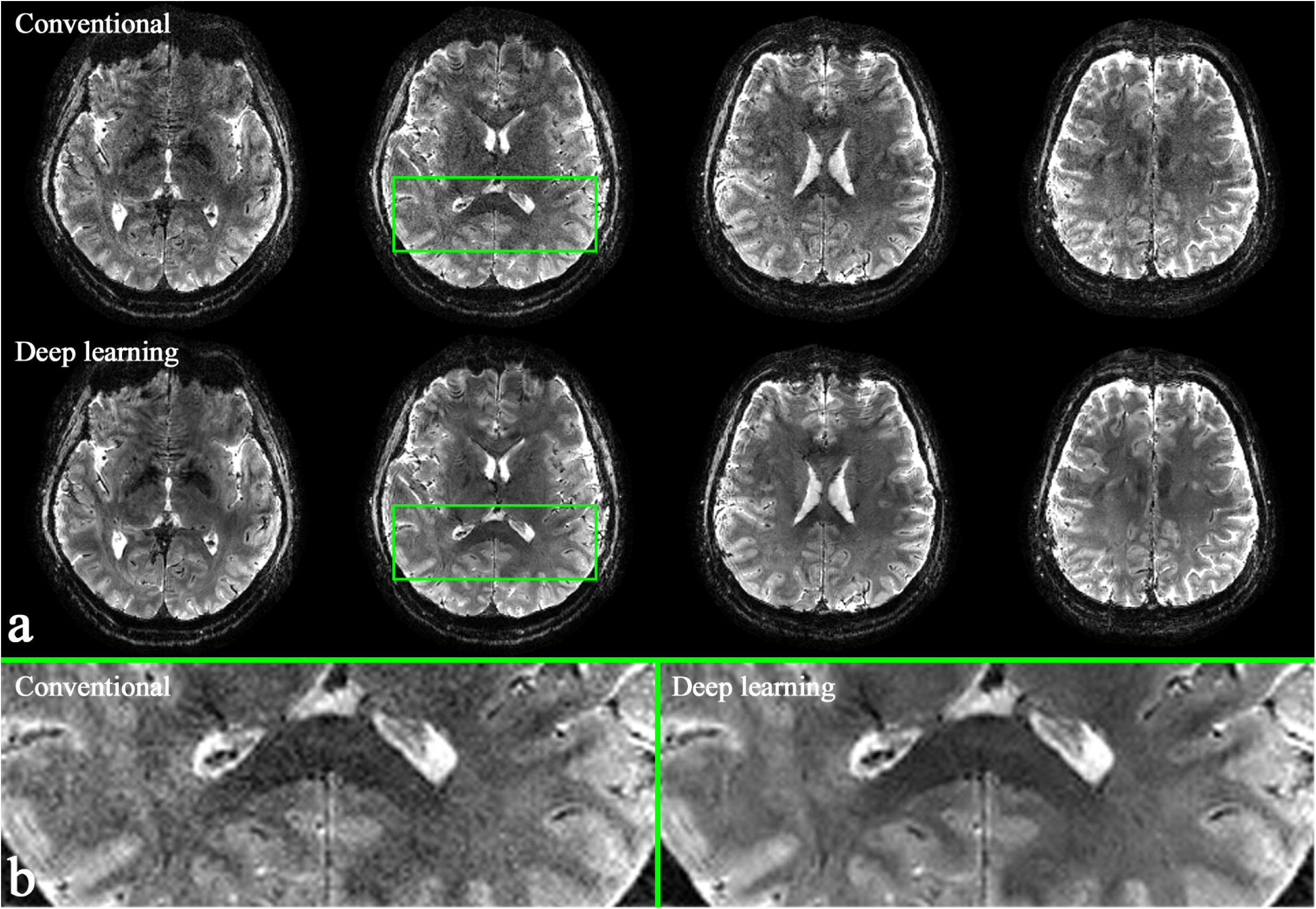
Reconstructed images from the submillimetre protocol (0.875□×□0.875□mm^2^). (**a**) Results obtained from the conventional (top row) and the deep learning (bottom row) methods at a representative slice location. (**b**) An enlarged depiction of the selected green ROIs from panel (a), highlighting the significant SNR improvement achieved with the deep learning technique.

**Figure 4.**
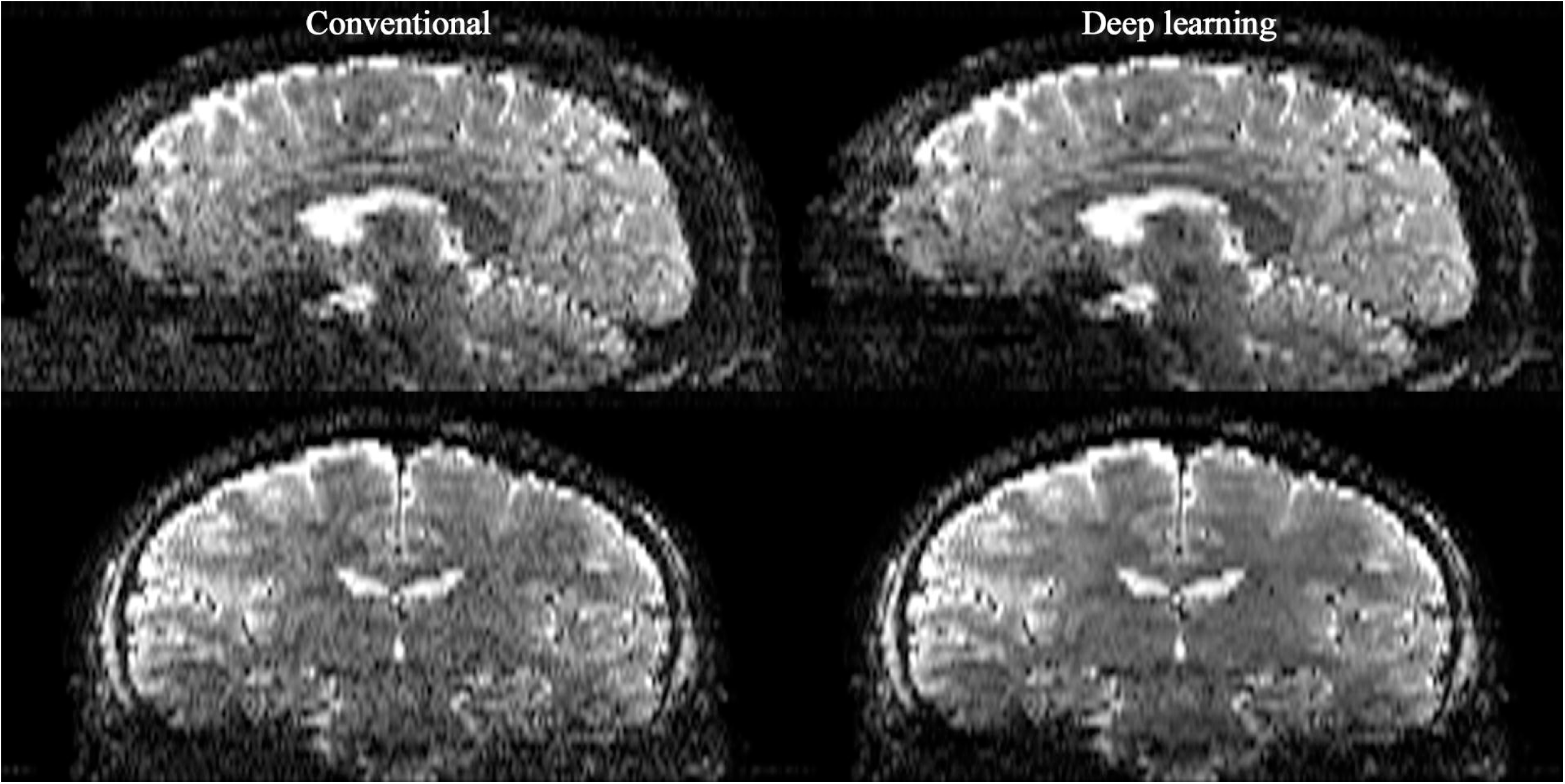
Results of resliced images in sagittal and coronal directions. The SNR enhancement in the deep-learning reconstructed images is evident across all slices, demonstrating the reliable performance of the developed method.

### 3.2. Histogram distribution

Figure 5a presents the deep learning-reconstructed image along with its corresponding GM and WM masks at a representative slice location. The histogram distribution of the signals within each ROI is shown in Fig. 5b, where the results from both the conventional and deep learning reconstruction methods are overlaid on the same plot for comparison. The dispersion of the histogram distribution was reduced in both ROI cases, with a more pronounced improvement in the WM, implying that signal variations caused by the noise contamination was substantially diminished. Table 1 presents the mean ± SD of signals computed for each ROI, demonstrating a reduction in signal standard deviation in the deep learning reconstruction results compared to the conventional method.

**Figure 5.**
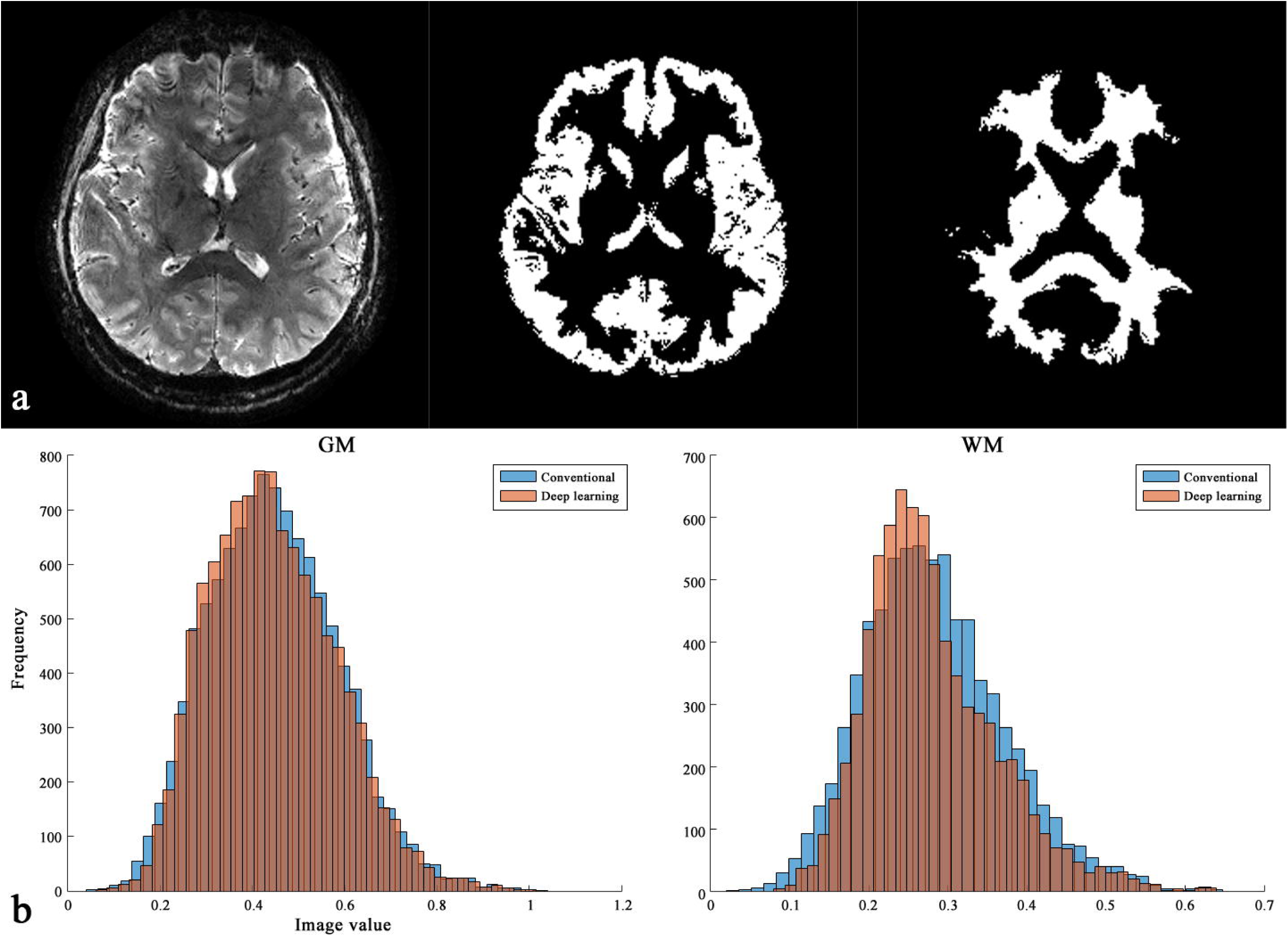
Analysis of histogram distribution. (**a**) A reconstructed slice from the deep learning technique with the corresponding GM and WM masks. (**b**) Histogram distributions of signals within GM (left) and WM (right) ROIs. Particularly for WM, the dispersion of the histogram distribution was significantly reduced, implying that signal variations caused by noise contamination was diminished.

**Table 1.** Comparison of mean ± SD of signals between the conventional and deep learning methods.

|  | GM | WM |
| --- | --- | --- |
| **Conventional** | $0.447 \pm 0.140$ | $0.283 \pm 0.089$ |
| **Deep learning** | $0.444 \pm 0.137$ | $0.279 \pm 0.081$ |

## 4. Conclusions and discussion

This study demonstrates the application of a deep learning network for the reconstruction of ultra-high-resolution fMRI data. The functional protocol was configured using the TR-external EPIK method, which has previously been shown to identify cortical depth-dependent functional profiles with a half-millimetre in-plane spatial resolution for various resting-state networks distributed throughout the brain from a single fMRI session (Yun et al., 2022). As our work represents the first application of deep learning reconstruction for TR-external EPIK, the spatial resolution and brain coverage used in the present work (0.875 × 0.875 mm^2^; 48 slices) was lower than in our early work (0.515 × 0.515 mm^2^; 108 slices), facilitating a more straightforward development of the deep learning network. The significantly smaller number of slices in this work was primarily due to the absence of multi-band acceleration. In future work, our network model will be further adapted for datasets with even higher spatial resolution and expanded brain coverage. Since our earlier study demonstrated reliable and reproducible detection of cortical depth-dependent functional profiles using only conventional reconstruction methods, the integration of deep learning techniques suggests the potential for TR-external EPIK to achieve spatial resolutions below half-millimetre. This advancement is expected to enable the exploration of layer-specific neural activities and underlying connectivity with greater precision and deeper insights.

A significant improvement in SNR was observed in the reconstructed images from deep learning reconstruction when compared those from the conventional method. This enhancement was consistent across all slice locations, demonstrating the reliability of the developed scan-specific deep-learning technique. The results of the histogram distribution of signals for the GM and WM regions further quantitatively confirm this enhancement, as the dispersion of the signal distribution is substantially reduced. This reduction was more pronounced for WM when compared to GM, which was mostly due to the spatially dependent differences in SNR characteristics of the employed RF receiver coil (32-channel phase-array coil), where signals are often higher near coil elements and lower in deeper regions. As a result, regions such as the centre of the brain (i.e. WM) experience greater signal degradation with lower SNR. Additionally, the coil geometry factor (g-factor) generates varying noise amplification across the in-plane dimension, further contributing to the reduced SNR in WM in the reconstructed images obtained from the conventional method. We adopted a self-supervised learning approach to train the network. For each slice, we generated 20 different sampling-split scenarios, yielding 960 unique batches per volume. For six time points, we had 5,760 batches. To ensure robust performance, a single network was trained across all slices and time points. With a batch size of two, training took roughly two hours per epoch for 48 slices and 6 time points, and we used 34 epochs in total. This training duration can be markedly reduced through transfer learning. Once trained, inference required around 40 seconds per time point.

Visual inspection of the reconstructed images suggests that the deep learning technique improved SNR without a significant loss of spatial resolution, which is beneficial to preserve original functional signals, ultimately contributing to the delineation of cortical depth-dependent profiles with enhanced accuracy. The current work focuses on the development of the deep learning network and the validation of its use with datasets acquired using ultra-high-resolution TR-external EPIK. In future studies, the technique will be applied to fMRI datasets to investigate putative advantages of the enhanced image reconstruction in detection of layer-specific functional activities. Additionally, the impact of this improvement on functional analysis will be thoroughly evaluated in comparison to existing de-noising techniques developed for the same purpose.

Notwithstanding the significant SNR improvement demonstrated in this work, the deep learning technique also holds potential for simultaneously correcting other image artefacts, such as aliasing. However, this improvement was not substantial with the employed datasets, as these artefacts were not evident in the conventional reconstruction method. The effectiveness of our method in reducing both noise and image artefacts will be demonstrated in future work by applying it to various datasets. Our deep learning method can be further improved by incorporating reversed phase-encoding data to address the geometric distortions typically seen in the EPI-based techniques. Although EPI techniques offer relatively high temporal resolution, enabling dynamic MRI studies such as fMRI, their high susceptibility to field inhomogeneities results in substantial geometric distortions in reconstructed images, subsequently hindering accurate mapping of neural activity onto anatomical references. Geometric distortion can be effectively mitigated using the reversed phase-encoding data, leveraging the fact that the distortion develops in the opposite direction. Further development of this approach could lead to significant advancements in layer-specific fMRI techniques and analysis, opening new potential for insights and applications.

